# Epigenome Editing Durability Varies Widely Across Cardiovascular Disease Target Genes

**DOI:** 10.1101/2023.05.17.541156

**Authors:** Madelynn N. Whittaker, Lauren C. Testa, Aidan Quigley, Ishaan Jindal, Saúl V. Cortez-Alvarado, Ping Qu, Yifan Yang, Mohamad-Gabriel Alameh, Kiran Musunuru, Xiao Wang

**Author notes:** **Corresponding Author** Xiao Wang, Ph.D., Perelman School of Medicine at the University of Pennsylvania, 3400 Civic Center Blvd, Bldg 421, 11-189 Smilow Center for Translational Research Philadelphia, PA 19104, USA.

## Abstract

**Background:** Hepatic knockdown of the proprotein convertase subtilisin/kexin type 9 (*PCSK9*) gene or the angiopoietin-like 3 (*ANGPTL3*) gene has been demonstrated to reduce blood low-density lipoprotein cholesterol (LDL-C) levels, and hepatic knockdown of the angiotensinogen (*AGT*) gene has been demonstrated to reduce blood pressure. Genome editing can productively target each of these three genes in hepatocytes in the liver, offering the possibility of durable “one-and-done” therapies for hypercholesterolemia and hypertension. However, concerns around making permanent gene sequence changes via DNA strand breaks might hinder acceptance of these therapies. Epigenome editing offers an alternative approach to gene inactivation, via silencing of gene expression by methylation of the promoter region, but the long-term durability of epigenome editing remains to be established.

**Methods:** We assessed the ability of epigenome editing to durably reduce the expression of the human *PCSK9, ANGPTL3*, and *AGT* genes in HuH-7 hepatoma cells. Using the CRISPRoff epigenome editor, we identified guide RNAs that produced efficient gene knockdown immediately after transfection. We assessed the durability of gene expression and methylation changes through serial cell passages.

**Results:** Cells treated with CRISPRoff and *PCSK9* guide RNAs were maintained for up to 124 cell doublings and demonstrated durable knockdown of gene expression and increased CpG dinucleotide methylation in the promoter, exon 1, and intron 1 regions. In contrast, cells treated with CRISPRoff and *ANGPTL3* guide RNAs experienced only transient knockdown of gene expression. Cells treated with CRISPRoff and *AGT* guide RNAs also experienced transient knockdown of gene expression; although initially there was increased CpG methylation throughout the early part of the gene, this methylation was geographically heterogeneous—transient in the promoter, and stable in intron 1.

**Conclusions:** This work demonstrates precise and durable gene regulation via methylation, supporting a new therapeutic approach for protection against cardiovascular disease via knockdown of genes such as *PCSK9*. However, the durability of knockdown with methylation changes is not generalizable across target genes, likely limiting the therapeutic potential of epigenome editing compared to other modalities.

## INTRODUCTION

Atherosclerotic cardiovascular disease (ASCVD) is the leading cause of death worldwide. Among the most firmly established causal risk factors for ASCVD are elevated blood lipid factors—namely, low-density lipoprotein cholesterol (LDL-C) and triglyceride-rich-lipoproteins— and elevated blood pressure. Proprotein convertase subtilisin/kexin type 9 (PCSK9), an antagonist to the LDL receptor, angiopoietin-like 3 (ANGPTL3), an inhibitor of lipoprotein lipase and endothelial lipase, and angiotensinogen (AGT), a component of the renin-angiotensin-aldosterone system, have all emerged as attractive therapeutic targets for the prevention of ASCVD. Naturally occurring loss-of-function variants in *PCSK9* or *ANGPTL3* cause reduced blood LDL-C levels and, in the case of *ANGPTL3*, reduced blood triglyceride-rich-lipoproteins; these variants confer substantial protection against ASCVD without evidence of serious adverse effects.^1–5^ Currently approved PCSK9 and ANGPTL3 inhibitors, whether monoclonal antibodies (evolocumab, alirocumab, and evinacumab) or short interfering RNA (inclisiran), are administered via subcutaneous or intravenous injection, and their effects are short-lived, ranging from a few weeks to several months.^6–9^ A short interfering RNA inhibiting AGT (zilebesiran) is currently being evaluated in clinical trials, with injections administered every several months.^10^

Hepatic genome editing to inactivate *PCSK9, ANGPTL3*, or *AGT* at the DNA level, by introducing loss-of-function variants, has been demonstrated in preclinical animal models including non-human primates and rats.^11–14^ Base-editing strategies targeting *PCSK9* or *ANGPTL3* have proven to be particularly effective and have demonstrated favorable off-target profiles thus far. Nonetheless, concerns about the permanence and irreversibility of DNA-level changes might limit the acceptance of genome-editing therapies for lipid or blood pressure modification. Durable yet potentially reversible inactivation of *PCSK9, ANGPTL3*, or *AGT* without the need to alter the DNA sequence could prove much more attractive to providers and patients.

In contrast to standard nuclease and base editors, epigenome editors utilize a catalytically dead Cas9 (dCas9) protein with a standard guide RNA (gRNA), maintaining the canonical search-and-bind functionality of nuclease editors without instigating double-strand DNA breaks or single-strand nicks. Epigenome editors modulate the epigenome via histone modifications, DNA methylation, and other changes, and when targeted to a gene regulatory unit, can substantially modify expression of the target gene. Recently, a set of CRISPR-based epigenome editing tools termed CRISPRoff and CRISPRon were reported to regulate gene expression via site-directed methylation and demethylation of gene promoters, respectively.^15^ CRISPRoff exerts its suppressive effects by methylating DNA in promoter/enhancer regions. This methylation inhibits the binding of transcription factors and recruits proteins capable of remodeling chromatin, which assumes the compact and transcriptionally inactive form of heterochromatin (**Figure 1**). As opposed to epigenome editors that rely on steric hindrance via dCas9 to repress transcription and transiently silence gene expression (CRISPR interference), CRISPRoff enables heritable and potentially more long-term gene silencing via methylation.^15^

**Figure 1.**
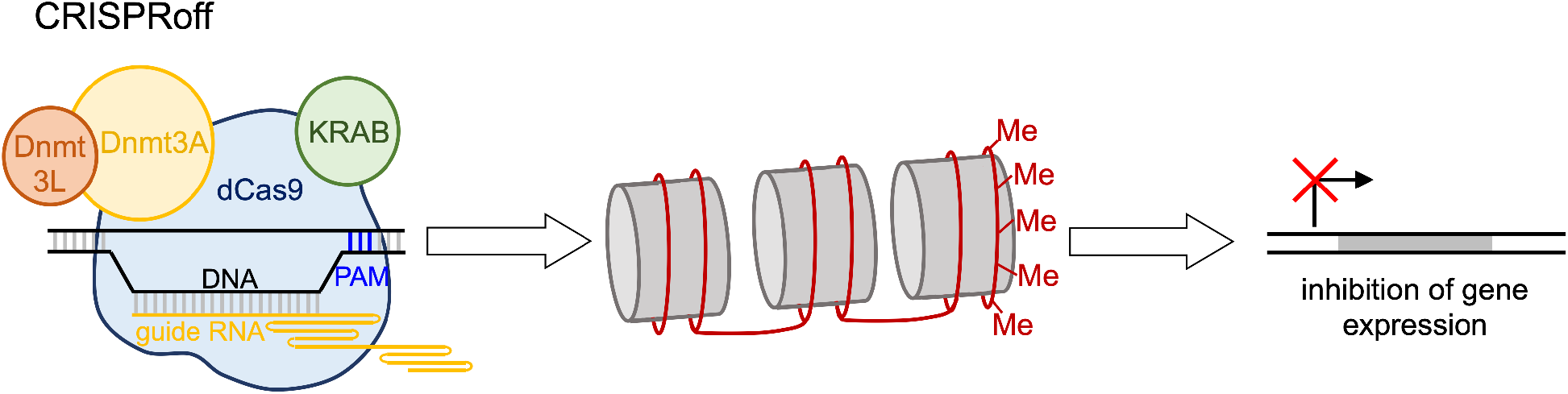
Schematic of CRISPRoff epigenome editing. dCas9 (catalytically dead Cas9) in complex with a guide RNA (gRNA) targets a desired genomic site proximal to a PAM (protospacer-adjacent motif) in the promoter of a gene. The KRAB (Krüppel associated box) repressor domain engages with chromatin at the site while Dnmt3a, facilitated by Dnmt3L, adds Me (methyl) groups on cytosine bases in the surrounding DNA sequence, particularly cytosine bases in CpG dinucleotide sequences. Methylation of a gene regulatory unit has the effect of reducing gene expression.

We hypothesized that CRISPRoff could durably induce methylation changes in the *PCSK9, ANGPTL3*, and *AGT* promoters and thereby modulate their expression. To demonstrate the viability of a possibly safer alternative to permanent gene knockout, epigenome editing was used to silence each of these three genes in HuH-7 hepatocyte-like cells with the goal of sustained methylation of the promoter and reduction of gene expression levels.

## RESULTS

### Epigenome editing of *PCSK9*

We first screened CRISPRoff with gRNAs targeting the *PCSK9* promoter, individually and in combinations, for their ability to reduce *PCSK9* expression in the human HuH-7 hepatoma cell line (**Figure 2A**). We focused on three CpG sites in the *PCSK9* promoter that were annotated in the UCSC Genome Browser as being unmethylated in hepatoma cells but methylated in other cell types (spanning hg38/chr1:55039312-55039339), suggesting a potential causal role in the preferential expression of *PCSK9* in hepatocytes. We evaluated 19 individual gRNAs (**Supplemental Table 1**), binding within 100 base pairs of these three CpG sites, by co-transfecting a plasmid expressing CRISPRoff and a plasmid expressing the gRNA into HuH-7 cells. We observed substantial suppression of *PCSK9* mRNA levels measured three days after transfection (**Figure 2A**). The average reduction in *PCSK9* mRNA expression for these gRNAs was 59%, with a maximum reduction of 79% (with P13 gRNA).

**Figure 2.**
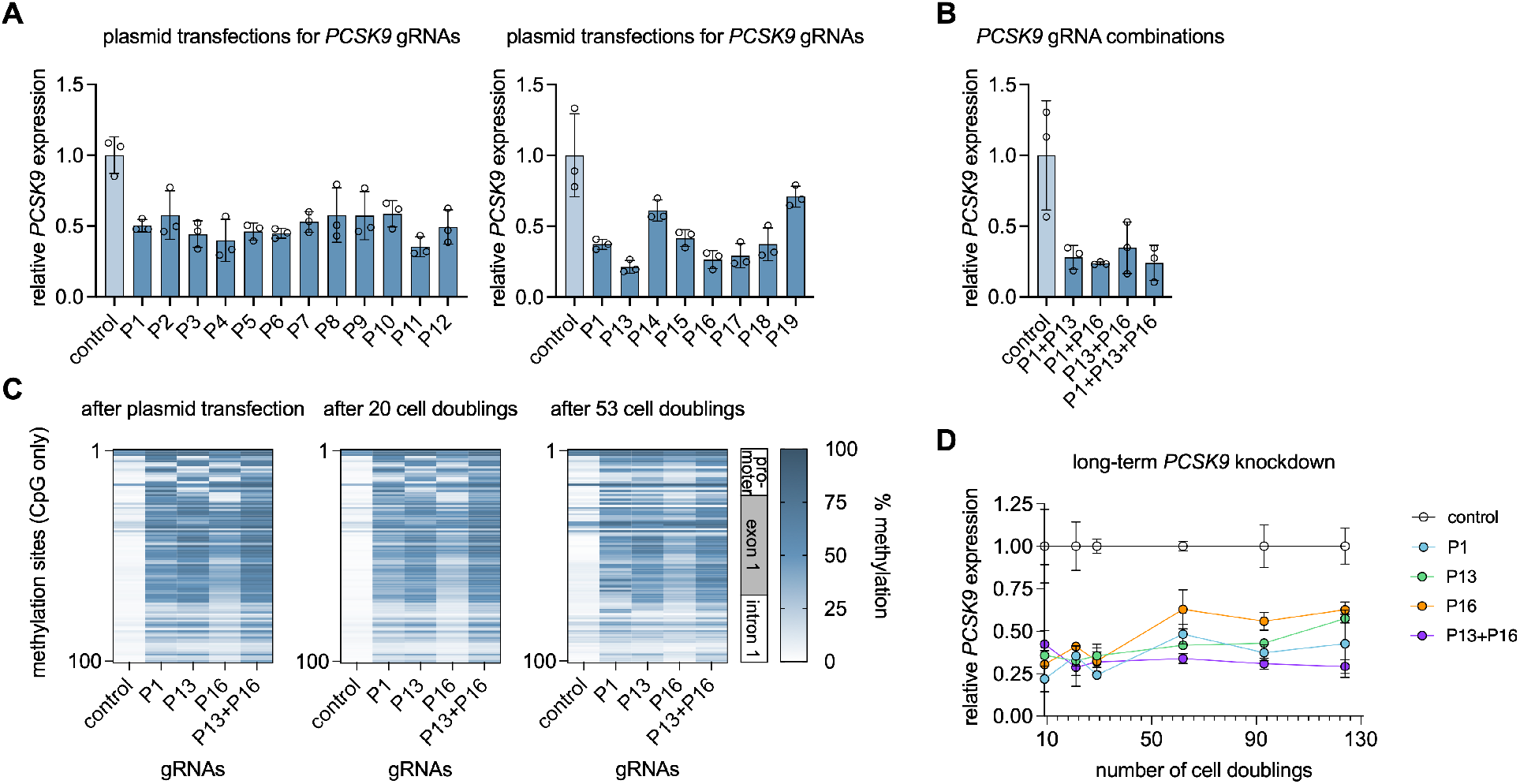
Epigenome editing of *PCSK9*. **A** and **B** show the effects of transfections with CRISPRoff plasmid and any of 19 gRNA plasmids, combinations of gRNA plasmids, or an empty gRNA plasmid (control), on *PCSK9* expression in HuH-7 cells three days after transfection, normalized to expression in control cells. Data are displayed as means and standard deviations (*n* = 3 biological replicates). **C** shows methylation in the early part of the *PCSK9* gene in HuH-7 cells at various times after plasmid transfection. **D** shows long-term effects on *PCSK9* expression in HuH-7 cells (same passaged cells used in **C**) at various times after plasmid transfection, with expression levels normalized to those of contemporaneous control cells. Data are displayed as means and standard deviations (*n* = 3 biological replicates).

We investigated the efficacy of four combinations of the best performing individual gRNAs: P1+P13, P1+P16, P13+P16, and P1+P13+P16 (**Figure 2B**). All combinations showed comparable efficacy to each other and to the individual gRNAs. We proceeded with P13+P16 because each gRNA has a very high MIT specificity score (99 for each gRNA, **Supplemental Table 1**), presumptively limiting off-target epigenome editing.

We then performed long-term experiments with the lead individual or combined gRNAs (P1, P13, P16, and P13+P16), assessing for both *PCSK9* gene expression and DNA methylation patterns in the *PCSK9* promoter. We found that the CRISPRoff gRNAs all induced profound increases in methylation at CpG dinucleotides in the *PCSK9* promoter (**Figure 2C**), with up to ≈80% decreases in *PCSK9* expression (**Figure 2D**). Notably, the individual gRNAs induced somewhat different methylation patterns throughout the *PCSK9* promoter, particularly in the vicinity of the gRNA target sites. These methylation increases and gene expression decreases endured through 124 cell doublings (i.e., 124 mitoses on average), with only mild attenuation over time. On average, the individual gRNAs initially had 67% knockdown of *PCSK9*, and after 124 cell doublings, the knockdown was 55%. In contrast, the P13+P16 combination proved to be more durable than any individual gRNA, as its ≈70% knockdown of *PCSK9* persisted through 124 cell doublings.

### Epigenome editing of *ANGPTL3*

In parallel, we tested the ability of CRISPRoff to silence the expression of *ANGPTL3*. As with *PCSK9*, we screened gRNAs targeting the promoter of the *ANGPTL3* gene and assessed both gene expression and methylation. We selected 6 gRNAs broadly distributed throughout the *ANGPTL3* promoter and with high MIT specificity scores (**Supplemental Table 1**) to test via plasmid transfection in HuH-7 cells and observed substantial suppression of *ANGPTL3* mRNA levels measured three days after transfection (**Figure 3A**). The mean reduction in ANGPTL3 mRNA expression for these gRNAs was 41% with a maximum reduction of 56%. After identifying the lead individual gRNAs (A1, A3, A4, A6) and various combinations of these gRNAs, we evaluated *ANGPTL3* mRNA knockdown and methylation of the *ANGPTL3* promoter. The dual gRNA knockdown approach, on average, reduced *ANGPTL3* expression by 73%, with a maximum reduction of 80% (A1+A4) (**Figure 3B**). We also observed substantial increases in methylation at some sites in the *ANGPTL3* promoter, despite the dearth of CpG dinucleotides in the promoter; methylation was increased on cytosine bases in non-CpG dinucleotides, particularly CpA dinucleotides (**Figure 3C**).

**Figure 3.**
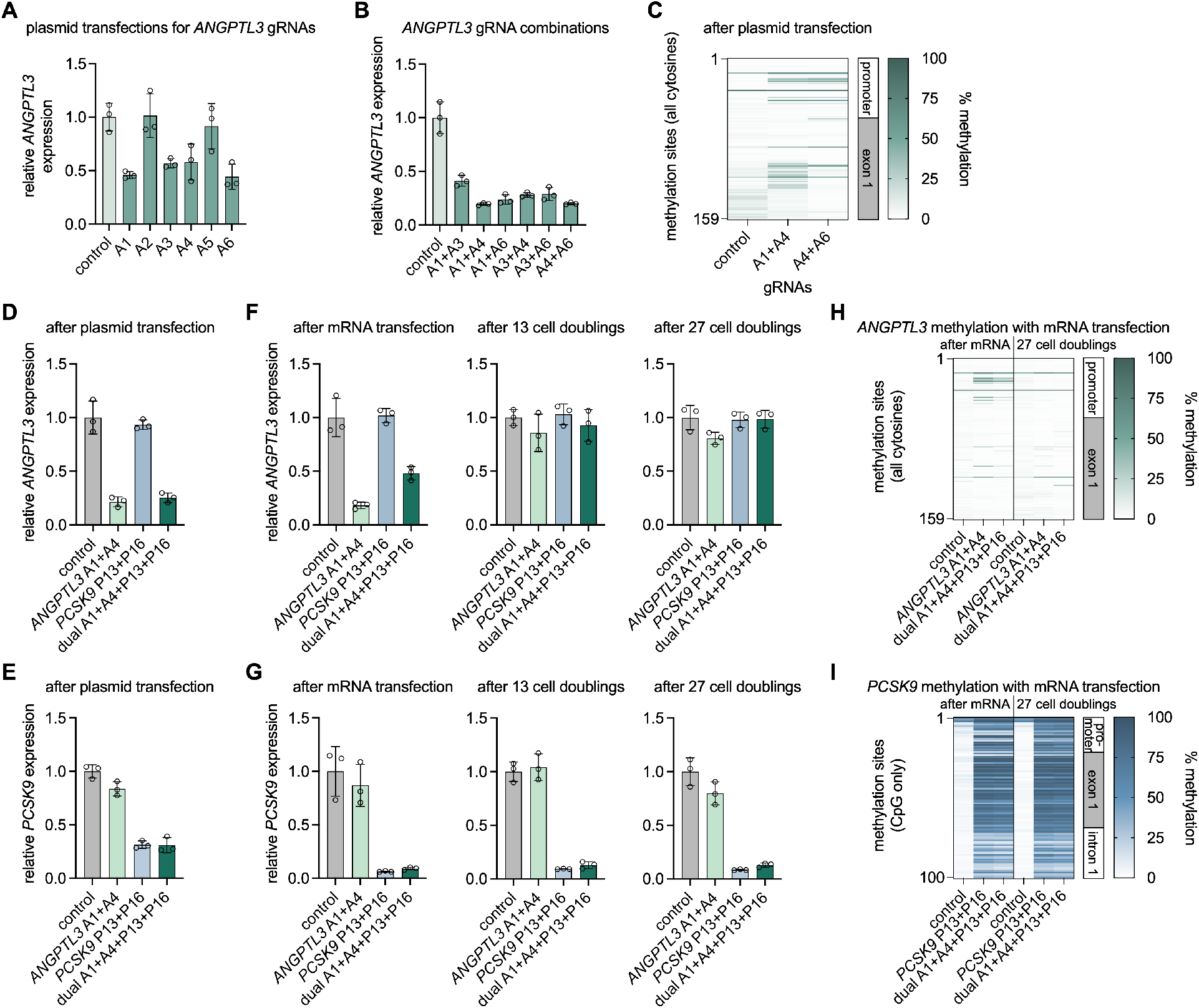
Epigenome editing of *ANGPTL3*. **A** and **B** show the effects of transfections with CRISPRoff plasmid and any of 6 gRNA plasmids, combinations of gRNA plasmids, or an empty gRNA plasmid (control), on *ANGPTL3* expression in HuH-7 cells three days after transfection, normalized to expression in control cells. Data are displayed as means and standard deviations (*n* = 3 biological replicates). **C** shows methylation in the early part of the *ANGPTL3* gene in HuH-7 cells three days after plasmid transfection. **D** and **E** show the effects of transfections with CRISPRoff plasmid and two *ANGPTL3* gRNA plasmids, two *PCSK9* gRNA plasmids, all four gRNA plasmids, or an empty gRNA plasmid (control), on (**D**) *ANGPTL3* expression and (**E**) *PCSK9* expression in HuH-7 cells three days after transfection, normalized to expression in control cells. Data are displayed as means and standard deviations (*n* = 3 biological replicates). **F** and **G** show the effects of transfections with CRISPRoff mRNA and synthetic gRNAs, paralleling the experiment in **D** and **E**, on (**F**) *ANGPTL3* expression and (**G**) *PCSK9* expression in HuH-7 cells at various times after transfection, normalized to expression in control cells. Data are displayed as means and standard deviations (*n* = 3 biological replicates). **H** and **I** show methylation in the early part of (**H**) the *ANGPTL3* gene or (**I**) the *PCSK9* gene in HuH-7 cells (same passaged cells used in **F** and **G**) at various times after mRNA transfection.

We then tested the ability to use a combination of gRNAs targeting *ANGPTL3* and *PCSK9* to simultaneously silence the expression of the two genes. With plasmid co-transfection, we introduced CRISPRoff and various combinations of gRNAs into HuH-7 cells: *ANGPTL3* alone (A1+A4), *PCSK9* alone (P13+P16), and both *PCSK9* and *ANGPTL3* (multiplex; A1+A4+P13+P16). After three days, the *ANGPTL3* gRNAs alone and the full multiplex combination both achieved 75-80% reduction of *ANGPTL3* expression, whereas the *PCSK9* gRNAs alone did not alter *ANGPTL3* expression (**Figure 3D**). In contrast, the *PCSK9* gRNAs alone and the full multiplex combination of four gRNAs both achieved ≈70% reduction of *PCSK9* expression, whereas the *ANGPTL3* gRNAs alone did not alter *PCSK9* expression (**Figure 3E**).

After establishing robust short-term knockdown of *ANGPTL3* and *PCSK9* using plasmid transfection, we adopted a different transfection methodology to validate the epigenome editing. Using *in vitro* transcribed CRISPRoff mRNA, we co-transfected mRNA and various combinations of synthetic gRNAs in HuH-7 cells. After three days, we observed substantial knockdown of *ANGPTL3* and *PCSK9* surpassing that achieved by plasmid transfection, ≈80% for *ANGPTL3* and ≈95% for *PCSK9* (**Figure 3F, 3G**). However, by 13 cell doublings, the knockdown effect on *ANGPTL3* had largely reversed, whereas the knockdown effect on *PCSK9* was stable and persisted through 27 cell doublings. Consistent with these observations, methylation in the *ANGPTL3* promoter was increased at three days but had largely reverted to baseline by 27 cell doublings, whereas methylation in the *PCSK9* promoter was profoundly increased at three days and remained similarly elevated at 27 cell doublings (**Figure 3H, 3I**).

### Epigenome editing of *AGT*

We next tested the ability of CRISPRoff to silence the expression of *AGT*. Unlike the promoter of *ANGPTL3*, the promoter of *AGT* harbors a number of CpG dinucleotides. We selected 8 gRNAs broadly distributed throughout the *AGT* promoter and with high MIT specificity scores (**Supplemental Table 1**) to test via plasmid transfection in HuH-7 cells and, as with *PCSK9* and *ANGPTL3*, observed substantial suppression of *AGT* mRNA levels measured three days after transfection, as much as 63% (**Figure 4A**). We then evaluated the combination of the two best performing gRNAs (AG5+AG8) via co-transfection of CRISPRoff mRNA and synthetic gRNAs. After three days, there was 92% knockdown of *AGT* expression (**Figure 4B**). However, unlike *PCSK9* and like *ANGPTL3*, by 14 cell doublings the knockdown effect on *AGT* had largely reversed. We observed greatly increased methylation of CpG dinucleotides in the *AGT* promoter, exon 1, and intron 1 at three days after transfection (**Figure 4C**). Intriguingly, by 14 cell doublings, the methylation in the promoter and exon 1 had largely reverted to baseline, whereas the methylation in intron 1 was stable, indicating that (1) the HuH-7 cells had spontaneously reverted CpG methylation in a geographically constrained manner following epigenome editing and (2) *AGT* expression correlates to *AGT* promoter methylation.

**Figure 4.**
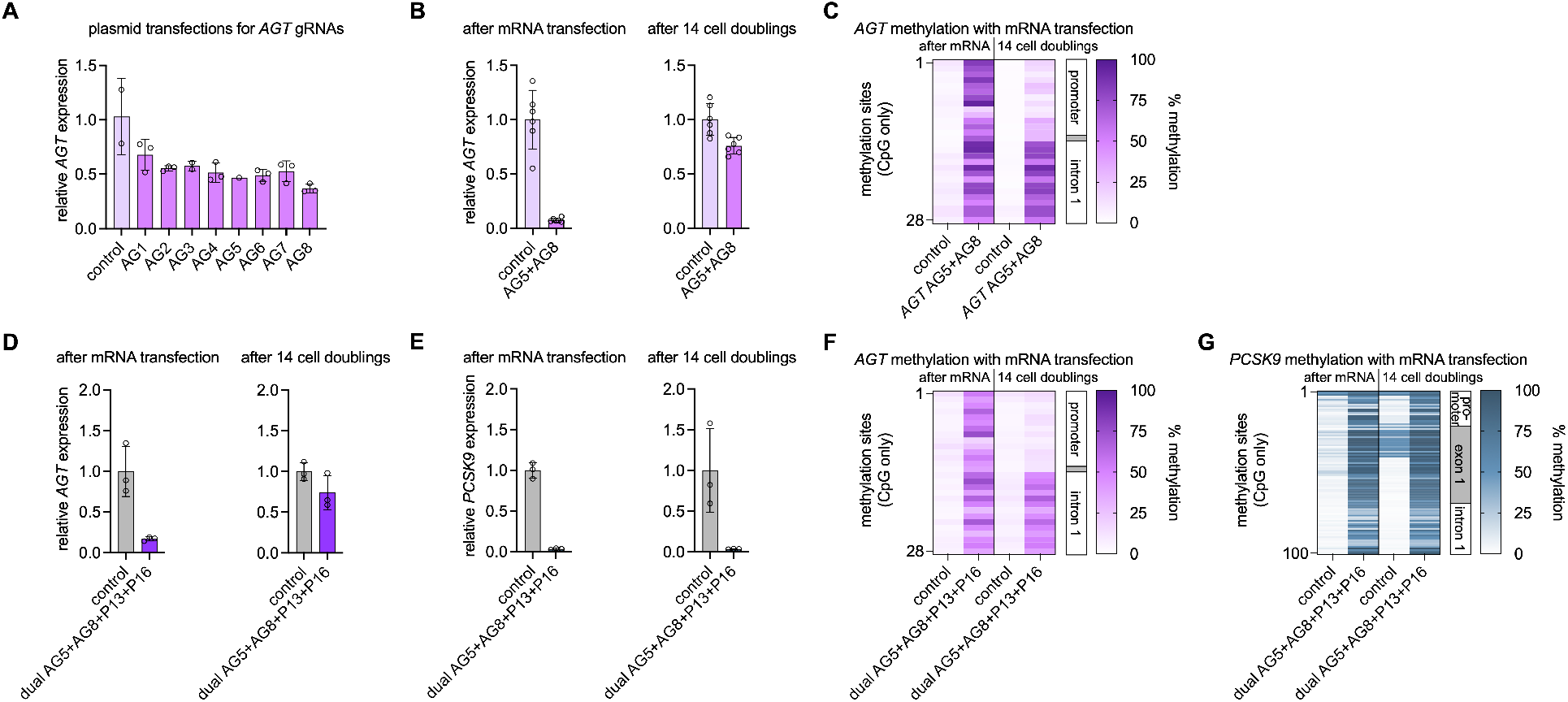
Epigenome editing of *AGT*. **A** shows the effects of transfections with CRISPRoff plasmid and any of 8 gRNA plasmids, or an empty gRNA plasmid (control), on *AGT* expression in HuH-7 cells three days after transfection, normalized to expression in control cells. Data are displayed as means and standard deviations (*n* = up to 3 biological replicates). **B** shows the effects of transfections with CRISPRoff mRNA and two synthetic gRNAs on *AGT* expression in HuH-7 cells at various times after transfection, normalized to expression in control cells. Data are displayed as means and standard deviations (*n* = 6 biological replicates). **C** shows methylation in the early part of the *AGT* gene in HuH-7 cells (same passaged cells used in **B**) at various times after mRNA transfection. **D** and **E** show the effects of transfections with CRISPRoff mRNA and a combination of four synthetic gRNAs (two for *AGT*, two for *PCSK9*) on (**D**) *AGT* expression and (**E**) *PCSK9* expression in HuH-7 cells at various times after transfection, normalized to expression in control cells. Data are displayed as means and standard deviations (*n* = 3 biological replicates). **F** and **G** show methylation in the early part of (**F**) the *AGT* gene or (**G**) the *PCSK9* gene in HuH-7 cells (same passaged cells used in **D** and **E**) at various times after mRNA transfection.

We then co-transfected mRNA and a multiplex combination of synthetic gRNAs targeting both *AGT* and *PCSK9* (AG5+AG8+P13+P16) into HuH-7 cells. After three days, *AGT* expression had decreased by 83% and *PCSK9* expression by 97% in the cells (**Figure 4D, 4E**). At 14 cell doublings, the cells continued to have decreased *PCSK9* expression (down by 96%) but *AGT* expression had rebounded (down by only 26%). Analysis of methylation in these cells demonstrated stably increased methylation in *PCSK9*, whereas in *AGT* the increased methylation was stable only in intron 1 and was transient in the promoter and exon 1, as observed in the prior experiment (**Figure 4F, 4G**).

### Global transcriptomic profiling of epigenome editing of *PCSK9*

An important consideration for characterizing programmable therapeutic editing therapies is off-target editing. The extent to which epigenome editors like CRISPRoff affect the expression of genes other than the target gene remains to be fully clarified. To address this, we performed RNA sequencing on HuH-7 cells co-transfected with CRISPRoff mRNA and synthetic *PCSK9* P13+P16 gRNAs, after 27 cell doublings, to assess for genome-wide perturbation of gene expression. We observed that the *PCSK9* gene had by far the most significant difference between control cells and treated cells (**Figure 5**). There were a handful of other genes whose expression differences reached statistical significance. While it is formally possible that some of these genes reflect off-target effects of CRISPRoff activity, they might also represent genes that are secondarily affected by dysregulation of PCSK9 activity in the cells.

**Figure 5.**
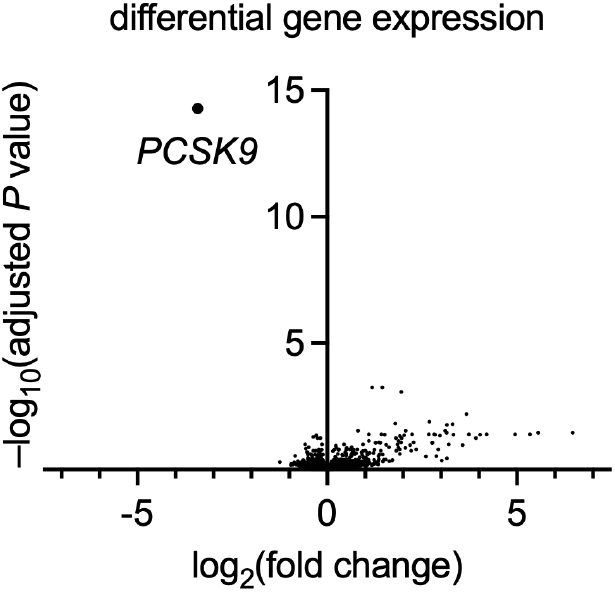
Differential gene expression with epigenome editing of *PCSK9*. Volcano plot of results from RNA sequencing of HuH-7 cells (same as passaged cells in **Figure 3F, 3G**) transfected with CRISPRoff mRNA and synthetic *PCSK9* P13+P16 gRNAs, compared with control cells, at 27 cell doublings after transfection (*n* = 3 biological replicates per group).

### DISCUSSION

Unlike other gene editing therapies—nuclease editing, base editing, prime editing, etc.^16^—a single dose of a CRISPRoff epigenome editing therapy could have a long-lasting effect without cleaving DNA strands and without changing the DNA sequence. Our study demonstrates that epigenome editing can achieve efficacy and durability of site-specific methylation with concomitant gene silencing, with the caveat that these effects are highly variable across loci. The three examples of therapeutic genes presented here are instructive with respect to the variability.

The early part of the *PCSK9* gene is rich in CpG dinucleotides (CpG island), and methylation of the cytosine bases by CRISPRoff was robust throughout the promoter and extended into exon 1 and intron 1. Despite the brief exposure to CRISPRoff, especially when delivered via mRNA, the methylation was durable through many cell doublings. This durability is consistent with the repeated action of DNA methyltransferases to enforce methylation of CpG dinucleotides on newly synthesized DNA strands templated on the parental strands already bearing methylation at those sites. The durable methylation was accompanied by durable, substantial reduction of *PCSK9* expression, up to 124 cell doublings over the course of several months in culture, especially with a combination of two gRNAs. Perturbation of *PCSK9* expression far exceeded the effects on any other genes in the genome. Given our observations in cultured hepatocyte-like cells, we anticipate that long-term durability, even lifelong durability, might manifest with the delivery of CRISPRoff and *PCSK9* gRNA into the liver in vivo in rodents, large animals, and perhaps ultimately in human beings.

The early part of the *ANGPTL3* gene has very few CpG dinucleotides. Although we observed methylation of cytosine bases in the short term, along with gene silencing, the methylation largely occurred in CpA dinucleotides. Given the lack of symmetry between the two DNA strands with CpA dinucleotides, and the lack of action of DNA methyltransferases at those sites, it was perhaps predictable that the methylation and gene silencing would not be durable, especially after a few rounds of DNA synthesis and mitosis.

The *AGT* gene proved to be the most unpredictable case. Although not harboring a CpG island, the *AGT* promoter does have a number of CpG dinucleotide sequences, and CRISPRoff induced robust methylation of cytosine bases throughout the promoter and into exon 1 and intron 1 in the short term, concomitant with substantial reduction of *AGT* expression. However, the methylation was only partially durable, with the *AGT* promoter and exon 1 losing much of the methylation, whereas the increased methylation was maintained in intron 1. The overall effect was a reversal of the gene silencing, suggesting that *AGT* expression is tied to the methylation status of the promoter. This pattern of *AGT* methylation and expression was also observed in cells that simultaneously underwent CRISPRoff-mediated knockdown of *AGT* and *PCSK9* and that demonstrated broad stability of *PCSK9* methylation, confirming locus-specific heterogeneity rather than a nonspecific genome-wide effect in these cells. Of note, the original report describing CRISPRoff indicated that epigenetic silencing of genes without CpG islands could be “metastable” in some cases, but no mechanistic exploration of this phenomenon at the level of methylation was undertaken.^15^ Here, the mechanism by which CRISPRoff-induced methylation of CpG dinucleotides is selectively reversed only in the *AGT* promoter is unclear. Prior reports suggest that *AGT* promoter methylation is labile and can be influenced via the binding of transcription factors such as C/EBP at specific sites in the promoter.^17,18^ We speculate that transcription factor binding sites present in the *AGT* promoter but not in intron 1 are responsible for the discrepant stability in CpG methylation. If so, multiplex CRISPRoff editing of *AGT* and the gene(s) encoding the responsible transcription factor(s) could in principle stabilize *AGT* gene silencing, although the pleiotropic effects on other targets of the transcription factor(s) would likely limit the therapeutic viability of this strategy.

In summary, we find that epigenome editing can effect precise and durable gene silencing via methylation, supporting a new therapeutic approach for protection against ASCVD via knockdown of genes such as *PCSK9*. However, the durability of the effects on methylation and gene expression is highly heterogenous across gene loci—suggesting that epigenome editing will only be useful for a subset of knockdown targets, determined on a case-by-case basis, which could limit its therapeutic potential compared to other gene editing approaches.

## METHODS

### Guide RNA design and preparation

Using the web tool CRISPOR,^19^ gRNA spacer sequences were selected to target sites in the *PCSK9, ANGPTL3*, and *AGT* promoters, prioritizing sequences with high MIT specificity scores (**Supplemental Table 1**). Each spacer’s sense and antisense oligonulceotide strands were annealed and ligated into a gRNA expression vector (pGuide, Addgene plasmid #64711). After transformations into competent *E. coli* cells (Takara Bio, catalog #636763), colonies were picked, cultured, and miniprepped using Mini Spin Columns (Epoch Life Science, catalog #1910250). DNA preparations were quantified using a NanoDrop spectrophotometer (Thermo Fisher Scientific). Synthetic gRNAs were obtained from Integrated DNA Technologies.

### mRNA synthesis

A plasmid DNA template containing a codon-optimized CRISPRoff coding sequence, with the BFP (blue fluorescent protein) sequence removed, and a 3’ polyadenylate sequence was linearized. An in vitro transcription reaction containing linearized DNA template, T7 RNA polymerase, NTPs, and cap analog was performed to produce mRNA containing N1-methylpseudouridine. After digestion of the DNA template with DNase I, the mRNA product underwent purification and buffer exchange, and the purity of the final mRNA product was assessed with spectrophotometry and capillary gel electrophoresis. Elimination of double-stranded RNA contaminants was assessed using dot blots and transfection into human dendritic cells. Endotoxin content was measured using a chromogenic Limulus amebocyte lysate (LAL) assay, which was negative.

### CRISPRoff plasmid transfection

HuH-7 cells were cultured in Dulbecco’s Modified Eagle Medium (containing 4 mM L-glutamine and 1 g/L glucose) supplemented with 10% fetal bovine serum and 1% penicillin/streptomycin. HuH-7 cells were seeded at 350,000 cells per well in 6-well plates, and transfections were performed 16-24 hours following seeding. Each well was treated with 1.25 µg CRISPRoff plasmid (Addgene plasmid #167981), 1.25 µg gRNA plasmid (described above), 0.25 µg GFP plasmid (pmaxGFP, Lonza), and 8.25 µL TransIT-X2 (Mirus, catalog #MIR2300) suspended in a total of 250 µL of Opti-MEM I Reduced Serum Medium (Thermo Fisher Scientific, catalog #31985062). The DNA/TransIT/Opti-MEM mixture was incubated for 20 minutes at room temperature and then added dropwise onto the cells. Cells underwent expansion starting 72 hours post transfection, with 1:8 or 1:16 splits for each passage (≈1 cell doubling per day).

### CRISPRoff mRNA transfection

HuH-7 cells were cultured, maintained, and seeded as stated above. mRNA transfections were performed 16-24 hours following seeding. In one tube, 7.5 µL Lipofectamine MessengerMax (Thermo Fisher Scientific, catalog #LMRNA008) was added to 125 µL of Opti-MEM and incubated for 10 minutes. In another tube, 1.25 µg CRISPRoff mRNA and 1.25 µg total gRNA (individual gRNA or equal mix by weight of multiple gRNAs) was suspended in a total of 125 µL of Opti-MEM. Diluted mRNA was added to each tube of diluted MessengerMAX, and the subsequent mRNA/MessengerMax/Opti-MEM mixture was incubated for 5 minutes at room temperature and then added dropwise onto the cells. Cells underwent expansion starting 72 hours post transfection, with 1:8 or 1:16 splits for each passage (≈1 cell doubling per day).

### Nucleic acid isolation and bisulfite conversion

Genomic DNA was harvested from post-transfection HuH-7 cells using the DNeasy Blood & Tissue Kit (Qiagen, catalog #69506). Similarly, total RNA was extracted using the RNeasy Mini Kit (Qiagen, catalog #74106). Both DNA and RNA levels were quantified using a Nanodrop spectrophotometer. Bisulfite conversion of DNA was conducted via the Bisulfite Conversion Kit (Qiagen, catalog #59104).

### Reverse transcription and quantitative polymerase chain reaction

Reverse transcription on extracted RNA was conducted using SuperScript II reverse transcriptase (Thermo Fisher Scientific, catalog #18064014) to yield cDNA. Quantitative polymerase chain reaction was performed to measure the mRNA expression levels of *PCSK9, ANGPTL3*, or *AGT* relative to an internal control, *B2M* (Beta-2-Microglobulin), using TaqMan Gene Expression Master Mix (Thermo Fisher Scientific, catalog #4369016), and the following probes from Thermo Fisher Scientific: *PCSK9*, Hs00545399_m1; *ANGPTL3*, Hs00205581_m1; *AGT*, Hs01586213_m1; and *B2M*, catalog #4326319E. Each 10 μL quantitative polymerase chain reaction contained 5 μL TaqMan Gene Expression Master Mix, 0.5 μL *B2M* probe, 0.5 μL probe targeting the gene of interest, and 4 μL cDNA (diluted 1:10 with water) and was performed in technical duplicates. Reactions were carried out on the QuantStudio 7 Flex System (Thermo Fisher Scientific). Expression levels were quantified by the 2^−ΔΔCt^ method.

### Next-generation sequencing (NGS)

PCR reactions were performed using NEB EpiMark Hot Start Taq DNA Polymerase (New England Biolabs, catalog #M0490L). The following program was used for all bisulfite-converted genomic DNA PCRs: 98°C, 20 seconds; 40× (98°C, 20 seconds, 57°C, 30 seconds, 72°C, 10 seconds); 72°C, 2 minutes (primers and covered cytosine positions are listed in **Supplemental Tables 2–7**). PCR products were visualized via QIAxcel capillary electrophoresis (Qiagen) and then purified and normalized via an NGS normalization 96 well kit (Norgen BioTex Corp, catalog #61900). A secondary barcoding PCR was conducted to add Illumina barcodes (Nextera XT Index Kit V2 Set A and/or Nextera XT Index Kit V2 Set D, catalog #FC-131-2001 and #FC-131-2004) using ≈15 ng of first-round PCR product as template and NEBNext Polymerase (New England Biolabs, catalog #M0541S). Final libraries were quantified via Qubit (Qiagen) and then sequenced on a MiSeq sequencer (Illumina).

### Bisulfite sequencing analysis

Methylation data was analyzed using CRISPResso2 (version 2.2),^20^ where methylation frequency was calculated as the cytosine percent composition at each of the candidate methylation sites (**Supplemental Tables 3, 5**, and **7**).

### RNA sequencing analysis

RNA integrity was assayed using the RNA ScreenTape on the Agilent TapeStation system. All samples had an RNA integrity number (RIN) greater than 7.0. Libraries were prepared and sequenced at Azenta Life Sciences on Illumina HiSeq 2000/2500 systems with paired-end, 150-bp read lengths with a target read depth of ≈30 million reads per sample.

Paired-ended RNA sequencing reads were aligned to the human reference genome (GRCh38.104) using STAR (version 2.7.9a).^21^ Fastqc (version 0.11.7) (https://www.bioinformatics.babraham.ac.uk/projects/fastqc/) was used to examine the quality of reads. Picard (https://broadinstitute.github.io/picard/) was used to mark the duplicates, compute the RNA sequencing metrics, and estimate the library complexity. We then computed the raw read counts using the Rsubread package (version 2.0.3)^22^ and normalized the counts into counts per million (CPM) using the cpm function from the edgeR package (version 3.1.4).^23^ The genes with 25% of samples having CPM <1 were considered as low-expressed and removed from further analysis. Differential gene expression analysis was performed by the Limma package (version 3.56.1).^24^ Namely, the Voom function was used to transform the data ready for linear regression; the lmFit function was used for fitting the linear model; the Log2FC and adjusted *P* values reported by the toptable function were used to generate the volcano plot.

## ACKNOWLEDGMENTS

None.

## ABBREVIATIONS

AGT: Angiotensinogen
ANGPTL3: angiopoietin-like 3
ASCVD: atherosclerotic cardiovascular disease
RNA: guide
gRNA: low-density lipoprotein cholesterol
LDL-C: proprotein convertase subtilisin/kexin type 9, PCSK9.

---

**Supplemental Table 1.**
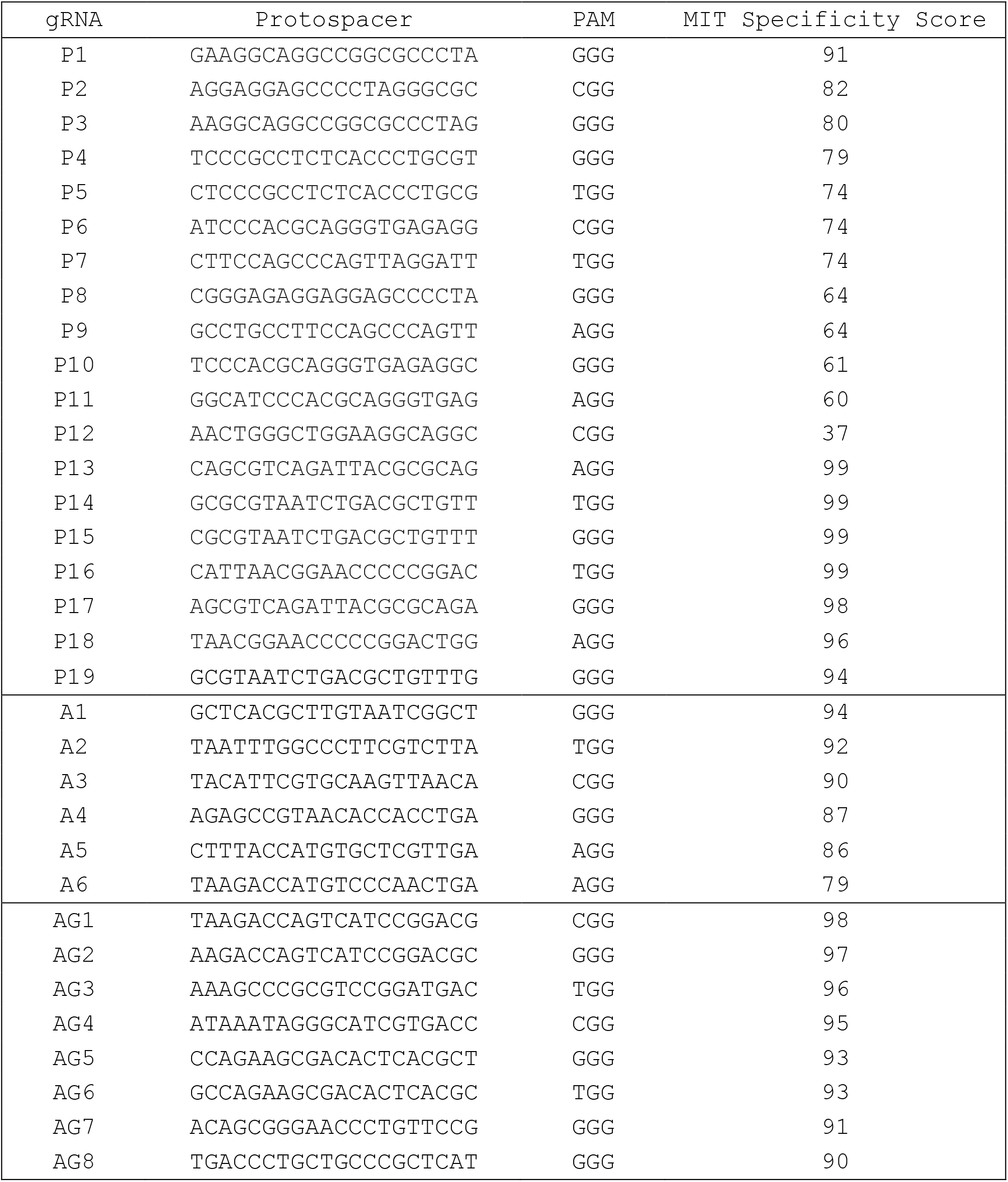
gRNAs used with CRISPRoff to target *PCSK9, ANGPTL3*, or *AGT*.

**Supplemental Table 2.**
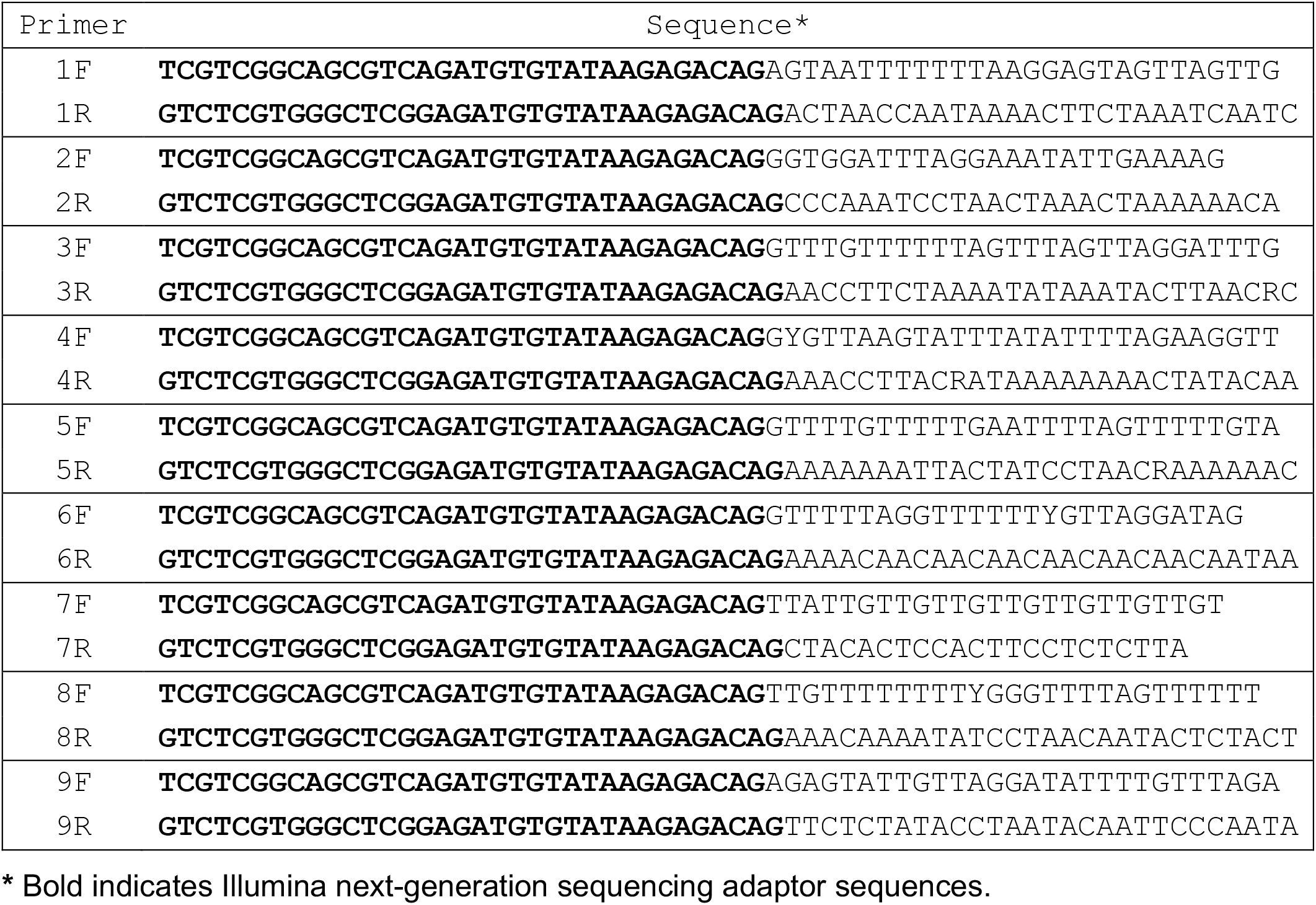
Primers for bisulfite sequencing of early part of *PCSK9*.

**Supplemental Table 3.**
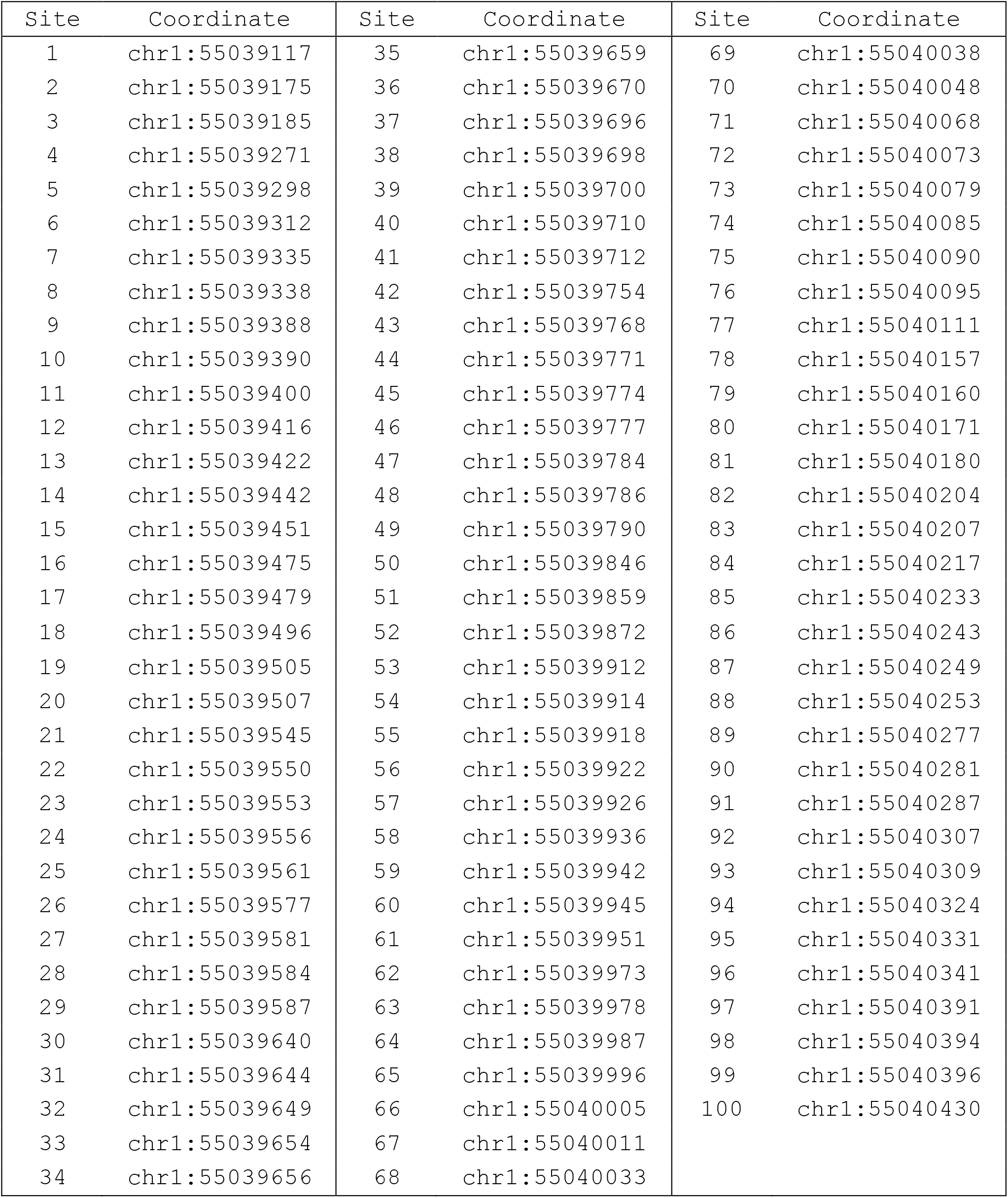
Genomic coordinates (hg38) of CpG sites in early part of *PCSK9*.

**Supplemental Table 4.**
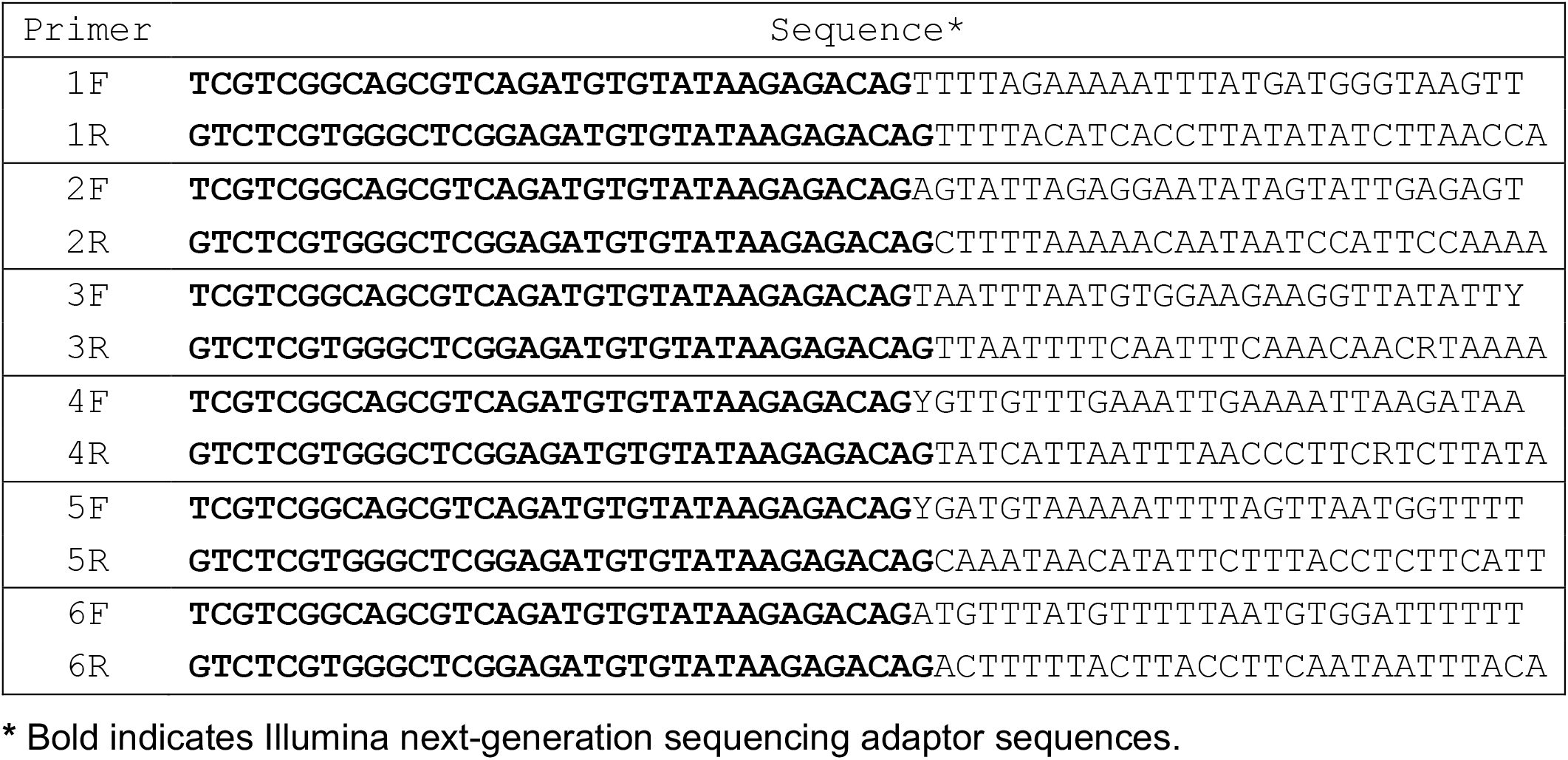
Primers for bisulfite sequencing of early part of *ANGPTL3*.

**Supplemental Table 5.** Genomic coordinates (hg38) of cytosine bases on sense strand in early part of *ANGPTL3*.

| Site | Coordinate | Site | Coordinate | Site | Coordinate |
| --- | --- | --- | --- | --- | --- |
| 1 | chr1:62596744 | 39 | chr1:62597418 | 77 | chr1:62597640 |
| 2 | chr1:62596751 | 40 | chr1:62597426 | 78 | chr1:62597649 |
| 3 | chr1:62596771 | 41 | chr1:62597428 | 79 | chr1:62597651 |
| 4 | chr1:62596776 | 42 | chr1:62597431 | 80 | chr1:62597655 |
| 5 | chr1:62596786 | 43 | chr1:62597443 | 81 | chr1:62597657 |
| 6 | chr1:62596787 | 44 | chr1:62597450 | 82 | chr1:62597658 |
| 7 | chr1:62596795 | 45 | chr1:62597452 | 83 | chr1:62597663 |
| 8 | chr1:62596798 | 46 | chr1:62597453 | 84 | chr1:62597664 |
| 9 | chr1:62596803 | 47 | chr1:62597456 | 85 | chr1:62597670 |
| 10 | chr1:62596804 | 48 | chr1:62597457 | 86 | chr1:62597679 |
| 11 | chr1:62596811 | 49 | chr1:62597460 | 87 | chr1:62597689 |
| 12 | chr1:62596812 | 50 | chr1:62597461 | 88 | chr1:62597706 |
| 13 | chr1:62596815 | 51 | chr1:62597465 | 89 | chr1:62597707 |
| 14 | chr1:62596818 | 52 | chr1:62597466 | 90 | chr1:62597713 |
| 15 | chr1:62596822 | 53 | chr1:62597471 | 91 | chr1:62597714 |
| 16 | chr1:62596824 | 54 | chr1:62597476 | 92 | chr1:62597716 |
| 17 | chr1:62596836 | 55 | chr1:62597506 | 93 | chr1:62597717 |
| 18 | chr1:62596837 | 56 | chr1:62597512 | 94 | chr1:62597720 |
| 19 | chr1:62596843 | 57 | chr1:62597515 | 95 | chr1:62597729 |
| 20 | chr1:62596845 | 58 | chr1:62597518 | 96 | chr1:62597735 |
| 21 | chr1:62596851 | 59 | chr1:62597519 | 97 | chr1:62597743 |
| 22 | chr1:62596853 | 60 | chr1:62597529 | 98 | chr1:62597749 |
| 23 | chr1:62596858 | 61 | chr1:62597572 | 99 | chr1:62597717 |
| 24 | chr1:62596860 | 62 | chr1:62597574 | 100 | chr1:62597720 |
| 25 | chr1:62596885 | 63 | chr1:62597582 | 101 | chr1:62597729 |
| 26 | chr1:62596894 | 64 | chr1:62597584 | 102 | chr1:62597735 |
| 27 | chr1:62596896 | 65 | chr1:62597585 | 103 | chr1:62597743 |
| 28 | chr1:62596900 | 66 | chr1:62597588 | 104 | chr1:62597749 |
| 29 | chr1:62596920 | 67 | chr1:62597600 | 105 | chr1:62597750 |
| 30 | chr1:62596927 | 68 | chr1:62597601 | 106 | chr1:62597757 |
| 31 | chr1:62597053 | 69 | chr1:62597603 | 107 | chr1:62597764 |
| 32 | chr1:62597058 | 70 | chr1:62597613 | 108 | chr1:62597765 |
| 33 | chr1:62597060 | 71 | chr1:62597614 | 109 | chr1:62597776 |
| 34 | chr1:62597061 | 72 | chr1:62597616 | 110 | chr1:62597783 |
| 35 | chr1:62597063 | 73 | chr1:62597617 | 111 | chr1:62597789 |
| 36 | chr1:62597064 | 74 | chr1:62597627 | 112 | chr1:62597791 |
| 37 | chr1:62597083 | 75 | chr1:62597632 | 113 | chr1:62597794 |
| 38 | chr1:62597086 | 76 | chr1:62597637 | 114 | chr1:62597804 |
| 115 | chr1:62597808 | 130 | chr1:62597736 | 145 | chr1:62597846 |
| 116 | chr1:62597819 | 131 | chr1:62597740 | 146 | chr1:62597847 |
| 117 | chr1:62597823 | 132 | chr1:62597741 | 147 | chr1:62597866 |
| 118 | chr1:62597825 | 133 | chr1:62597752 | 148 | chr1:62597868 |
| 119 | chr1:62597828 | 134 | chr1:62597760 | 149 | chr1:62597879 |
| 120 | chr1:62597832 | 135 | chr1:62597773 | 150 | chr1:62597897 |
| 121 | chr1:62597833 | 136 | chr1:62597782 | 151 | chr1:62597902 |
| 122 | chr1:62597842 | 137 | chr1:62597783 | 152 | chr1:62597915 |
| 123 | chr1:62597861 | 138 | chr1:62597796 | 153 | chr1:62597919 |
| 124 | chr1:62597871 | 139 | chr1:62597798 | 154 | chr1:62597925 |
| 125 | chr1:62597874 | 140 | chr1:62597799 | 155 | chr1:62597927 |
| 126 | chr1:62597882 | 141 | chr1:62597814 | 156 | chr1:62597933 |
| 127 | chr1:62597885 | 142 | chr1:62597815 | 157 | chr1:62597937 |
| 128 | chr1:62597890 | 143 | chr1:62597827 | 158 | chr1:62597938 |
| 129 | chr1:62597730 | 144 | chr1:62597845 | 159 | chr1:62597940 |

**Supplemental Table 6.**
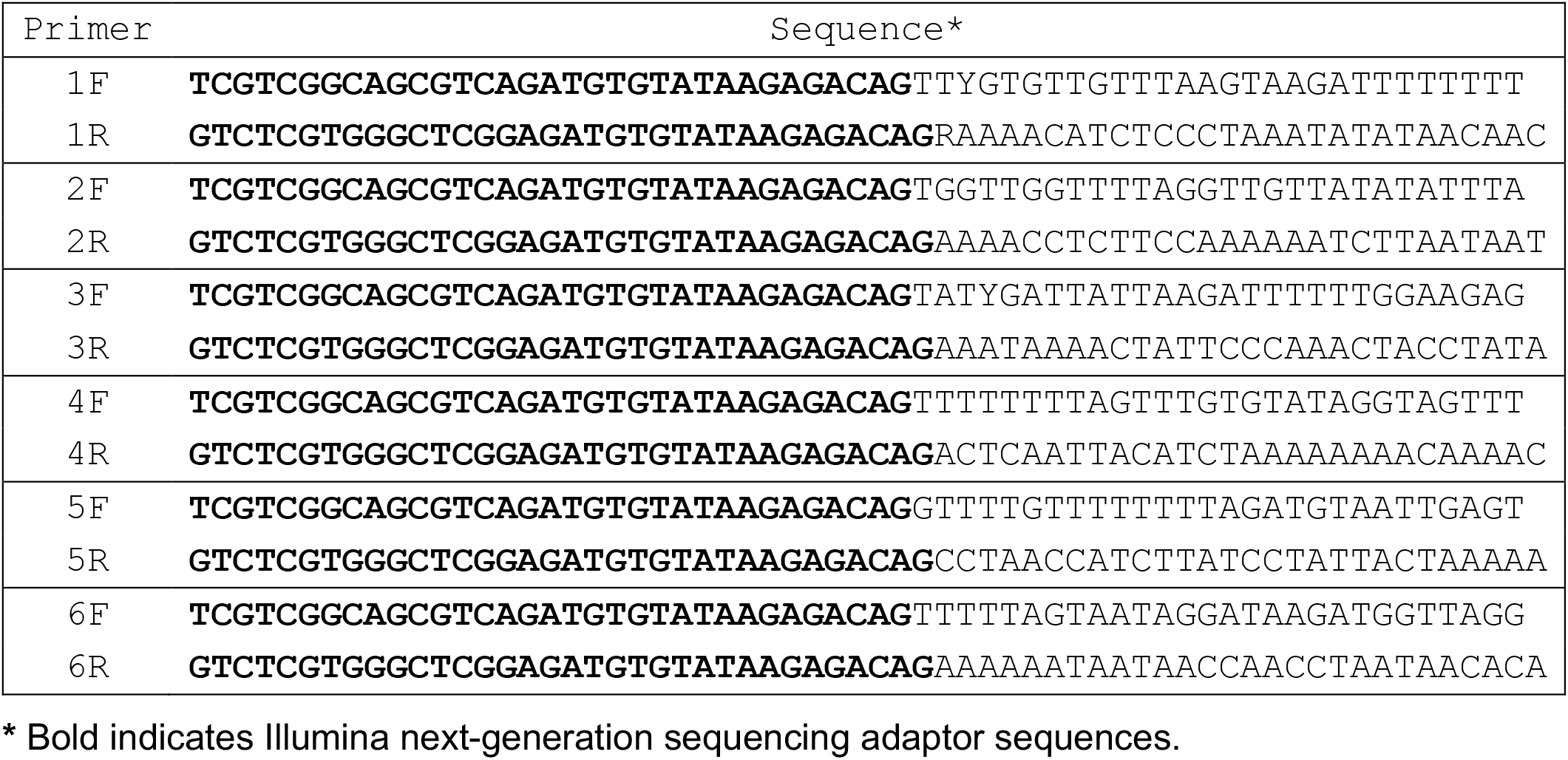
Primers for bisulfite sequencing of early part of *AGT*.

**Supplemental Table 7.**
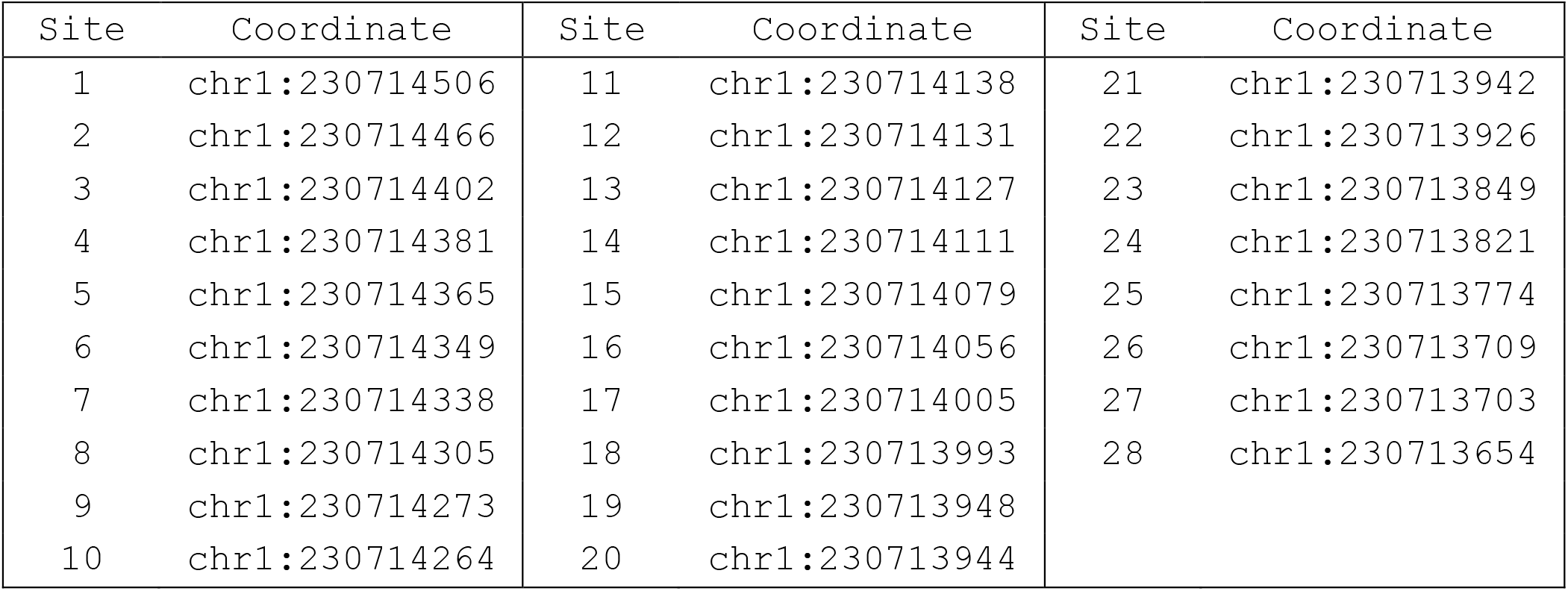
Genomic coordinates (hg38) of CpG sites in early part of *AGT*.

